# Taking off the training wheels: Measuring auditory P3 during outdoor cycling using an active wet EEG system

**DOI:** 10.1101/157941

**Authors:** Joanna E. M. Scanlon, Kimberley A. Townsend, Danielle L. Cormier, Jonathan W. P. Kuziek, Kyle E. Mathewson

## Abstract

Mobile EEG allows the investigation of brain activity in increasingly complex environments. In this study, EEG equipment was adapted for use and transportation in a backpack while cycling. Participants performed an auditory oddball task while cycling outside and sitting in an isolated chamber inside the lab. Cycling increased EEG noise and marginally diminished alpha amplitude. However, this increased noise did not influence the ability to measure reliable event related potentials (ERP). The P3 was similar in topography, and morphology when outside on the bike, with a lower amplitude in the outside cycling condition. There was only a minor decrease in the statistical power to measure reliable ERP effects. Unexpectedly, when biking outside significantly decreased P2 and increased N1 amplitude were observed when evoked by both standards and targets compared with sitting in the lab. This may be due to attentional processes filtering the overlapping sounds between the tones used and similar environmental frequencies. This study established methods for mobile recording of ERP signals. Future directions include investigating auditory P2 filtering inside the laboratory.

**Highlights:**

- A backpack containing all the equipment necessary to record ERP and EEG was worn by participants as they rode a bicycle outside along a street
- EEG and ERP data from an auditory oddball task is compared with data acquired within subject inside the lab
- Reliable MMN/N2b and P3 responses were measured during bicycle riding outside equal in magnitude to those obtained inside the lab
- A surprising decrease in the P2 component of the ERP evoked by targets and standards was observed when doing the task outside on a bicycle, which we attribute to increased auditory filtering

## 1. Introduction

Brain imaging has made significant advances in recent years, providing insight into the functional relationship between cognitive processes and brain activity (Makeig et al., 2010). However, the use of non-invasive technologies to record brain activity often requires participants to remain stationary (Debener et al., 2012). This is because natural sounds and sensations, as well as movement all have the potential to introduce noise into the EEG signal (Schlögl, Anderer, Roberts, Pregenzer, & Pfurtscheller, 1999; White & Van Cott, 2010), and this noise is typically what determines statistical power during EEG and ERP analysis (Luck, 2014). In efforts to begin expanding the limits of cognitive neuroscience recordings to mobile, real life settings, recent advances towards mobile electroencephalography (EEG) systems have provided a platform from which researchers can begin to study and understand the cognitive processes involved in critical human behaviours such as driving, exercise, and human interaction. The present study was conducted to attempt to overcome the limitations of measuring EEG on a moving participant, while determining the way in which cycling outside may influence auditory event related potentials (ERPs).

Recent progress in mobile EEG technology provides the possibility for significant developments in cognitive neuroscience. The majority of these studies make use of the oddball task due to this task’s high signal to noise ratio when detecting the P3, as well as for the ability to infer the attentional effects of a concurrent task (Polich, 1987; Polich & Kok., 1995). Debener et al. (2012) collected data for an oddball task completed sitting indoors or walking outdoors while brain activity was measured using a small consumer wireless EEG mounted to a classic laboratory electrode cap. The results were analyzed using brain-computer interface (BCI) single-trial P3 classification with linear discrimination. This analysis revealed that both the indoor (77%) and outdoor (69%) conditions had high prediction accuracies and the P3 generated indoors was significantly larger than the P3 found in the outside condition. However the authors noted that identifying whether the differences in P3 amplitude were a matter of added noise or distinct cognitive processes requires further research (Debener et al. 2012). The study aimed to reduce noise and signal degradation through the use of a small, wireless EEG system and concluded EEG data could be recorded while mobile in less than ideal conditions. While the authors showed that mobile EEG recordings are possible, the data collected did not achieve as high a quality as can be obtained from medical grade recording devices inside the lab. Additionally, Scanlon and colleagues (2017) collected EEG data for an auditory P3 oddball task during sub-aerobic indoor stationary cycling and sitting in a laboratory environment. The authors found that stationary cycling added noise to the data during and after pedaling, however reliable ERP signals could be accurately recorded, with no differences between sitting and cycling conditions. These studies demonstrate that mobile EEG recording is feasible, yet requires further study to achieve the same quality as data collected within the laboratory.

Many mobile EEG technologies stem from work with brain-computer interfaces (BCI), which allows one to control a computer using the classification of signals from the brain, such as ERPs and oscillations (de Vos et al., 2014). Using the same wireless mobile EEG system as Debener and colleagues (2012), de Vos and colleauges (2014) compared outdoor walking and outdoor sitting conditions while participants performed a three-stimulus oddball paradigm. This follow-up study accounted for possible outdoor distractions that could potentially influence task performance. The noise was not significantly different between conditions and single trial classification was above chance for both sitting and walking. The P3 elicited in response to targets was 35% smaller in the walking condition (De Vos et al., 2014). These results suggest that the differences in P3 are not a result of noise but instead some aspect of cognitive processing that was different between the two conditions. Gramann and colleagues (2010) purposed that P3 was smaller in the walking condition because cognitive resources were divided amongst large amounts of incoming muscular and sensory information. In another study using a passive electrode wireless mobile EEG system, Zink, Hunyadi, Van Huffel, and de Vos (2016) had participants perform an auditory oddball paradigm, while either sitting, pedaling, or cycling in an outdoor natural environment. They found no differences in RMS data noise or P3 between pedaling and sitting, but observed a nearly significant P3 amplitude decrease and increased RMS at outer electrode sites while participants were moving around. Additionally, BCI classification accuracies were significantly lower for the moving condition compared to the sitting still and pedaling conditions. While the current study does not use BCI classification, the purpose of the present study is to determine if EEG data collected outdoors using the current active electrode system can be of sufficient quality to be used as an input for BCI, potentially allowing for its use in natural environments.

In addition to ERPs, some studies have also looked at cortical oscillations during movement. Storzer and colleagues (2016) investigated oscillations in the motor cortex during stationary cycling and walking tasks of comparable speed. The authors demonstrated that while participants were cycling there were decreases in the high beta band (23-35 Hz) while movements were being initiated and executed, with subsequent increases in this band while movements were terminated. Additionally, both walking and cycling were associated with a decrease in alpha (8-12Hz) power with a significantly larger decrease during walking. Jain, Gourab, Schindler-Ivens, and Schmit (2013) also measured motor cortex oscillations using EEG on a stationary bicycle, demonstrating significant increases in beta desynchronization during active pedaling compared to passive (low-effort) pedaling. These studies indicate that it is possible to measure cortical oscillations during movement tasks, and that cycling may have effects on alpha and beta band oscillations.

Several studies have investigated ways in which to optimize EEG technologies to be better used in mobile situations. Makeig and colleagues (2010) described numerous features that are required for an effective mobile EEG system. These include EEG sensors that are small and lightweight to not hinder movement, removing the use of conductive gels to prevent electrical activity gathered from adjacent electrodes from combining together and finally, reducing the use of wires by implementing battery powered amplifiers and wireless data recording. Similarly, de Vos and Debener (2014) recommended that the mobile EEG technologies be small, lightweight and able to avoid cable motion (ideally wireless). In accordance with these recommendations, Debener, Emkes, Gandras, de Vos and Bleichner (2015) demonstrated that they could reliably record a P3 ERP component using a cEEGrid electrode array, made up of electrodes arranged in a flexible sheet and placed around the ear. A similar approach was used in a brain-computer interface (BCI) speller-task study by de Vos, Kroesen, Emkes, and Debener (2014), which showed equivalent results between a wired laboratory EEG system and a mobile wireless amplifier. Another study using a BCI speller by Bleichner and colleagues (2015) used a BCI system which could obtain significant P3 effects in a spelling task, while being small enough to hide under a baseball cap. The current study does not use many of the recommendations for an ideal mobile EEG system, and instead investigates the feasibility of using a regular (non-mobile) EEG system for a mobile task outside of the lab, as well as the ways in which the resulting data collected will differ from that collected within the typical laboratory environment.

An important consideration in the process of mobilizing our regular EEG system is the type of electrodes used, as this is the first source of data collection and can affect data quality in every stage of analysis. Several studies have investigated which electrode types might be best for non-ideal recording conditions such as mobile EEG in natural environments. Mathewson and colleagues (2017) compared active-dry electrodes to passive-wet and active-wet electrodes with the same amplifier. The electrodes were compared on single trial noise, event-related potentials, scalp topographies, ERP noise and ERP statistical power. The results reflected that active-dry electrodes had comparatively higher levels of noise than the passive-wet and active-wet systems, indicating that wet electrodes result in more statistical power and require less trials to get the same quality of data as dry electrode systems. Oliviera and colleagues (2016) investigated different electrode types in mobile studies by comparing BioSemi (active amplification) wet electrodes to Cognionics dry and wet electrodes (passive electrodes with active shielding) during an oddball task while participants were either walking or sitting. The BioSemi active wet electrodes showed the best results, with no difference in data noise between seated and walking conditions, while both passive dry and passive wet electrodes demonstrated significantly increased pre-stimulus noise and P3 time window amplitude variance, as well as decreased signal-to-noise ratio. While it may be argued that this study could have been confounded by the electrode caps and amplifier types used, active electrodes appear to offer some benefit to mobile EEG that warrants further study. Further, Laszlo, Ruiz-Blondet, Khlaifian, Chu, and Jin (2014) investigated the difference between active and passive electrodes using the same amplifier and different levels of impedance. The authors showed that while passive electrodes had ideal results in extremely low-impedance conditions (< 2 kΩ), active electrodes had the best results for all impedance levels above this point. With increased electrical noise and impedance due to wire movements being a major concern when recording EEG during mobile activity, this study adds to evidence that active amplification and wet electrodes may be the ideal for mobile EEG studies. While future research is needed to indicate the best type of electrodes specifically for mobile EEG research, the current study uses active wet electrodes based on the evidence available.

Few previous studies have directly measured whether accurate and statistically reliable ERP data can be collected during outdoor cycling. The current study is an extension to the findings of Scanlon, Sieben, Holyk, and Mathewson (2017), using similar methods and analysis. It is of particular interest to examine the potential ERP component variations between indoor laboratory conditions and outdoor path cycling, as well as how these conditions differ in statistical noise and power. Each participant completed four 6 minute blocks of an auditory P3 task while both sub-aerobically cycling on an outdoor path and sitting inside a laboratory faraday cage. Active, low-impedance wet electrodes (*actiCAP*) were used. The noise levels and power spectra were analyzed, as well as ERP morphology, topography, and magnitude for P3, P2 and MMN/N2b. Our first hypothesis is that due to a larger degree of movement during recording, the outside cycling condition will show an increased amount of ERP and single-trial noise, leading to decreased statistical power. Our second hypothesis is that even with the effects of this movement, we will be able to accurately record ERPs in both the inside sitting and outside cycling conditions, with a high enough level of accuracy to detect possible differences between conditions. Furthermore, our third hypothesis is that in accordance with previous mobile EEG evidence, attention will be split between cycling and the oddball task, demonstrated by a decreased P3 component during the outdoor cycling condition.

## 2. Materials and Methods

### 2.1. Participants

A total of 12 members of the university community participated in the experiment (Mean age=22.9; Age range=31-20; 4 female), and each participant was compensated 20$. Participants all had normal or corrected-to-normal vision with no history of neurological problems. The experimental procedures were approved by the Internal Research Ethics Board of the University of Alberta.

### 2.2. Materials

During the ‘outside’ condition, participants selected one of two bicycles (Kona Mahuna), differing in only frame size (17 in or 19 in) based on participant’s height. Seat height was adjusted to a comfort level as indicated by the participant. The bicycles were equipped with a small mock-press button, fastened on the right handlebar. Resistance was kept constant for all participants at a set level of 2^nd^ gear in both the front and back in order to allow participants to pedal evenly and constantly throughout the trials at a sub-aerobic level.

In the ‘inside’ condition, participants were seated in front of a 1920 x 1080 pixel ViewPixx/EEG LED monitor. A fixation cross was presented using a windows 7 PC running Matlab with the Psychophysics toolbox. Video output was via an Asus Striker GTX760. On the right armrest of the participant's’ chair was a mock-press button identical to the one placed on the bicycle. In both conditions, data was recorded and observed using a Microsoft Surface Pro 3 running Brain Vision recorder (Brain Products), and powered by an Anker Astro Pro2 20000mAh Multi-Voltage External Battery. The mentioned technology was connected using a Vantec 4-Port USB 3.0 Hub.

A Raspberry Pi 2 model B computer, which was running version 3.18 of the Raspbian Wheezy operating system, using version 0.24.7 of OpenSesame software (Mathôt, Schreij, & Theeuwes, 2012), was used both to run the oddball task and to mark the data for ERP averaging (see Kuziek, Sheinh, & Mathewson, 2017 for validation study). Audio output was via a 900MHz quad-core ARM Cortex-A7 CPU connected through a 3.5mm audio connector.

Coincident in time with sound onset, 8-bit TTL pulses were sent to the amplifier by a parallel port connected to the General Purpose Input/Output (GPIO) pins to mark the data for ERP averaging. A start button was affixed to the raspberry pi to simplify the beginning of a trial. Participants wore a two-pocket backpack (Lululemon; Fig. 1A-B) which contained all of the recording and stimulus generating materials. When outside, participants rode along a 1.4 km shared use path on Saskatchewan Drive in Edmonton, Canada (Figure 1C-D).

### 2.3. Procedure

Each participant completed an auditory oddball task to measure their P3 response to target tones in both the ‘inside’ and ‘outside’ condition (Figure 1E). Each condition was counterbalanced across participants, and was separated by a ten minute break and impedance check. Both inside and outside, participants completed four blocks of 250 trials separated by a 30 second break for a total of 1000 trials. Each trial had a 1/5 likelihood of being a target trial. Each block began with a 10-second countdown to ensure the oddball task only took place when the participant was already pedaling steadily. Each trial began with a pre-tone interval between 1000 and 1500 ms, followed by the tone onset. The next trial began immediately after the tone offset, with participants responding to targets during the following pre-tone interval. The headphones played one of two different frequency tones (either 1500 or 1000 Hz; sampled at 44.1 kHz; two channel; 16-ms duration; 2-ms linear ramp up and down), with the rare target tone always being 1500 Hz. In both conditions, the participant’s task was to press the mock-press button with the index finger of their right hand when the rare tone was heard. During the outside cycling condition, participants were instructed to pedal slowly and evenly, and try to mostly stare forward while performing the oddball task, and to limit eye movements. In the inside condition, participants were instructed to sit still and stare at the fixation while performing the identical oddball task.

**Figure 1.**
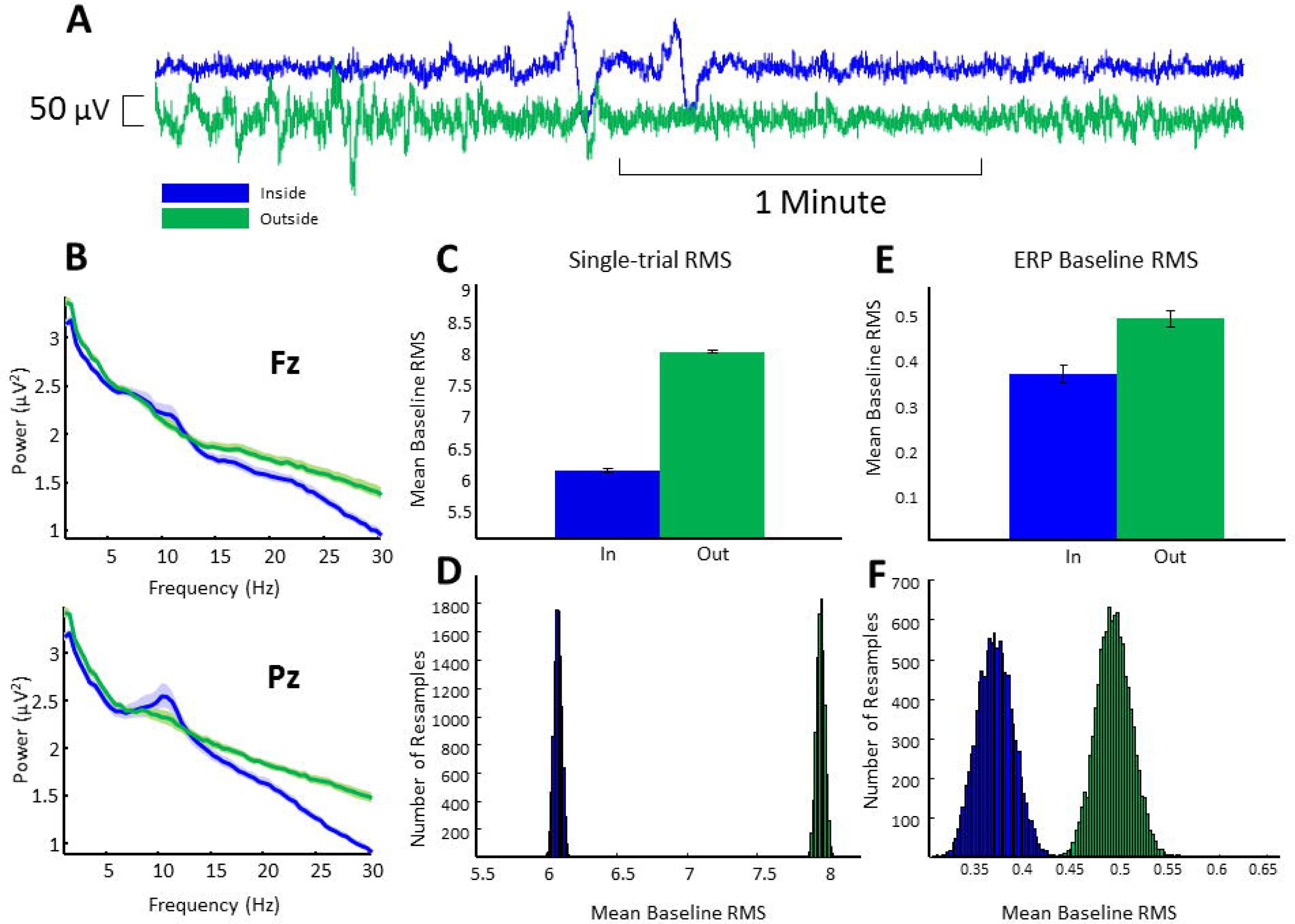
Mobile EEG biking apparatus and procedure. A: Participants performed the task while cycling on the bicycle, wearing the EEG cap and backpack containing the EEG equipment. B: Schematic layout of mobile EEG backpack contents. The two-pocket backpack contained the following: a Microsoft Surface Surface Pro 3, Anker Astro Pro2 20000mAh Multi-Voltage external battery, Raspberry Pi 2 model B, Brain Products v-Amp 16-channel amplifier, Vantec 4-Port USB 3.0 hub. C: The Procedure had two conditions: Biking outside and sitting inside. C: Shows the shared-use path participants rode on beside a 50 km/h two-way road. D: Shows the path on a map. E: Both sitting inside and outside on the bike, participants performed four 6 minute blocks of an auditory oddball tasks, with breaks in between. Within the oddball task, 80% of the tones were frequent low-pitched tones (1000Hz), while 20% were rare high-pitched tones (1500Hz), each played for 16 ms with a 2ms ramp-up and down. Tones were played 1-1.5 seconds apart.

### 2.4. EEG Recording

Based on previous lab work by Laszlo and colleagues’ (2014) comparing active and passive amplification electrodes at various levels of impedance, as well as a comparison by Oliviera and colleagues’ (2016) of active wet, passive wet, and passive dry electrodes during a mobile task, active wet electrodes (BrainProducts actiCAP) were selected for the study, as both of these studies demonstrated that they afford cleaner and better quality signals in non-ideal recording conditions. Ag/AgCl pin electrodes were used and arranged in 10-20 positions (Fp2, F3, Fz, F4, T7, C3, Cz, C4, T8, P7, P3, Pz, P4, P8, and Oz). Additionally, a ground electrode, embedded in the cap at position Fpz, and two reference electrodes, clipped to the left and right ear, were used. SuperVisc electrolyte gel and mild abrading of the skin with the blunted syringe tip were used to lower impedances of all the electrodes. Gel application and aforementioned techniques continued until impedances were lowered to < 10 kΩ, measured using an impedance measurement box (BrainProducts) and until data quality appeared clean and free of excessive noise. In addition to the 15 EEG sensors, 2 reference electrodes, and the ground electrode, the vertical and horizontal bipolar electrooculogram (EOG) was recorded from passive Ag/AgCl easycap disk electrodes affixed above and below the left eye, and 1 cm lateral from the outer canthus of each eye. Passive electrodes were used for the EOG as the AUX ports of our amplifier did not support active electrodes and we did not want to give up 4/16 EEG channels. These EOG channels can nonetheless record a reliable signal because of their placement on the face, where there is usually less hair to impede the signal, and any dirt and oils are removed before electrode placement. NuPrep preparation gel was applied to the applicable areas of the face, followed by wiping of the skin using an antibacterial sanitizing wipe, both used to lower the impedance of these EOG electrodes based on visual inspection of the data. These bipolar channels were recorded using the AUX ports of the V-amp amplifier, using a pair of BIP2AUX converters, and a separate ground electrode affixed to the central forehead.

EEG and EOG was recorded with a Brain Products V-amp 16-channel amplifier powered by the laptop USB port. Data were digitized at 500 Hz with a resolution of 24 bits. Data were filtered with an online bandpass with cutoffs of 0.1 and 30 Hz, along with a notch filter at 60 Hz. The narrow filters used were recommended in the actiCAP Xpress manual as ways to increase reliability and signal quality in mobile settings (Brain Products, 2014).

### 2.5. EEG Analysis

EEG analyses were computed using MATLAB 2012b with EEGLAB (Delorme & Makeig, 2004), as well as custom scripts. After recording, EEG data was re-referenced to the average of the left and right ear lobe electrodes. Timing of the TTL pulse was marked in the EEG data during recording, and used to construct 1200-ms epochs (including the 200-ms pretrial baseline) which were time-locked to the onset of standard and target tones. The average voltage in the first 200-ms baseline period was subtracted from the data for each electrode and trial. To remove artifacts caused by amplifier blocking, movement and any other non-physiological factors, any trials in either of the conditions with a voltage difference from baseline larger than ± 1000 *μ*V on any channel (including eye channels) were removed from further analysis. Over 98 percent of trials were retained after this procedure in each condition. This lenient threshold was chosen after also viewing all the analysis and figures with a second threshold after eye correct of ±500uV. Since the results were not different with this more strict rejection threshold we decided to show the results using only the 1000 uV threshold and eye correction. This allows us to keep as many trials for power analysis as possible, to allow approximately equal numbers of rejected trials for both conditions, and to get a sense of the true level of noise introduced by biking on the ERP measurement. On average, artifact rejection left approximately equal numbers of trials per participant in the cycling outside (M_targ_ = 199; range_targ_ =186-205; SD_targ_ = 5.0632; M_stand_ = 789; range_stand_ = 761-804; SD_stand_ = 14.7083), and sitting inside conditions (M_targ_ = 201; range_targ_ = 191-212; SD_targ_ = 4.7378; M_stand_ = 798; range_stand_ =788-806; SD_stand_= 4.7065; SD = Standard Deviation) from which we computed the remaining analyses.

A regression-based eye-movement correction procedure was used to estimate and remove variance due to artefacts in the EEG due to blinks, as well as vertical and horizontal eye movements (Gratton, Coles, & Donchin, 1983). This technique identifies blinks with a template-based approach, then computes propagation factors such as regression coefficients, predicting the horizontal and vertical eye channel data from the signals at each electrode. This eye channel data is then subtracted from each channel, weighted by these propagation factors, allowing us to remove most variance in the EEG which can be predicted by eye movements. No further rejection or filtering was done on the data in order to include as many trials as possible for both conditions, as well as to investigate how minor sources of non-eye-related noise contribute to the power to measure ERP components during the outdoor cycling task.

### 2.6. Condition Differences

When indoors, trials took place in a dimly lit sound and radio frequency attenuated chamber, with copper mesh covering the window. The only electrical devices in the chamber were an amplifier, speakers, keyboard, mouse, and monitor. The fan and DC lights were turned on, to allow proper ventilation and visual acuity of the fixation. The monitor runs on DC power from outside the chamber, the keyboard and mouse plugged into USB outside the chamber, and the speakers and amplifier were both powered from outside the chamber. Nothing was plugged into the internal power outlets. Any devices transmitting or receiving radio waves (i.e. cellphones) were either turned off or removed from the chamber for the duration of the experiment. The experiment was completed between the months of August and November. When outdoors, participants wore a toque over the electrodes to prevent the electrolyte gel used with the EEG cap from drying out, goggles to avoid added blinking during the trials, and gloves, if desired due to the weather (Temperature range = −5.3-10.4°C; Mean temperature =2.04°C). Each 6 minute block would bring the participant approximately 650 meters past the starting point, after which they would turn around, have the EEG impedance checked by a research assistant, and bike back.

## 3. Results

### 3.1. Data Noise

In Figure 2A raw data is depicted for a representative participant at the location of electrode Pz. To estimate the noise in the data on individual trials, we used two separate methods. First, we averaged the frequency spectra over each EEG epoch in the Pz and Fz electrode locations. Each participant’s data was sampled randomly for 504 of their artifact-removed standard trials. A fast Fourier transform (FFT) was then calculated by symmetrically padding the 600 time point epochs with zeros, making a 1,024-point time series for each epoch, providing frequency bins with a resolution of .488 Hz. Only frequencies measuring up to 30 Hz were plotted because data were collected online with a low-pass filter of 30Hz. Spectra for each participant were then calculated by averaging the 504 spectra from each participant, which were then combined into the grand-averaged spectra, demonstrated in Figure 2B for the Pz and Fz channels. Shaded regions indicate the standard error of the mean across participants. Evident from the graph is an increase in beta oscillations (15-30 Hz) during the outside biking compared to inside condition at both Pz (M _In-Out power_= −0.3272; SD_power_ = 0.1633; p = 2.4468e-05) and Fz (M _In-Out power_ = −0.2433; SD_power_ = 0.1734; p = 5.0314e-04) electrode locations, as tested using a two-tailed paired samples *t*-test. This difference was greatest in posterior locations, likely due to muscle artifacts in the neck during biking. Also evident from the graphs is an increase in delta (1-4Hz) frequencies during the outside condition at both Pz (M _In-Out power_ = −0.1247; SD_power_ = 0.1233; p = 0.0049) and Fz (M _In-Out power_ = −0.1127; SD_power_ = 0.1104; p = 0.0046).

Both conditions showed the expected 1/frequency structure in the data, while the inside condition demonstrated the typical peak in the alpha frequency range between 8 and 12 Hz over posterior locations (Mathewson et al., 2011). However, data from the outdoor cycling condition did not appear to indicate this peak. The outside condition was shown through a two-tailed paired samples *t*-test to have marginally lower power in alpha oscillations compared to the inside condition (M _In-Out power_ = 0.1209; SD_power_ = 0.2506; p = 0.1228).

### 3.2. Single-Trial Noise

As an additional estimate of the noise on single-trial EEG epochs, we calculated the RMS value of a baseline period for each trial (de Vos & Debener, 2014). This baseline period consisted of the 100 time points (200 ms) prior to each tone’s onset, to avoid including any interference of the evoked ERP activity in the measurement of RMS. RMS is equivalent to the average absolute voltage difference around the baseline, and therefore is a good estimate of EEG data single-trial noise. To estimate a distribution of RMS for each condition in our data, we used a permutation test that, without replacement, selects a different set of 360 epochs for each participant on each of 10,000 permutations before running second-order statistics (Laszlo et al., 2014; Mathewson et al., 2017). For each of these random selections and each condition, a grand average single-trial RMS was computed and recorded. A bar graph of the mean and standard deviation of grand average and single-trial RMS permutation distributions is shown in Figure 2C. Figure 2D shows a histogram of the grand-averaged single-trial RMS values calculated for each permutation, for each condition. Evident from these plots is a clear distinction between the single-trial noise between the two conditions. The outside condition (M_RMS-EEG_ = 7.9512; SD_RMS-EEG_ = 0.0291) clearly showed larger single-trial noise levels compared to the inside condition (M_RMS-EEG_= 6.0825; SD_RMS-EEG_ = 0.0276; Wilcoxon rank sum test; z = −122.4714; p < 0.0001).

**Figure 2.**
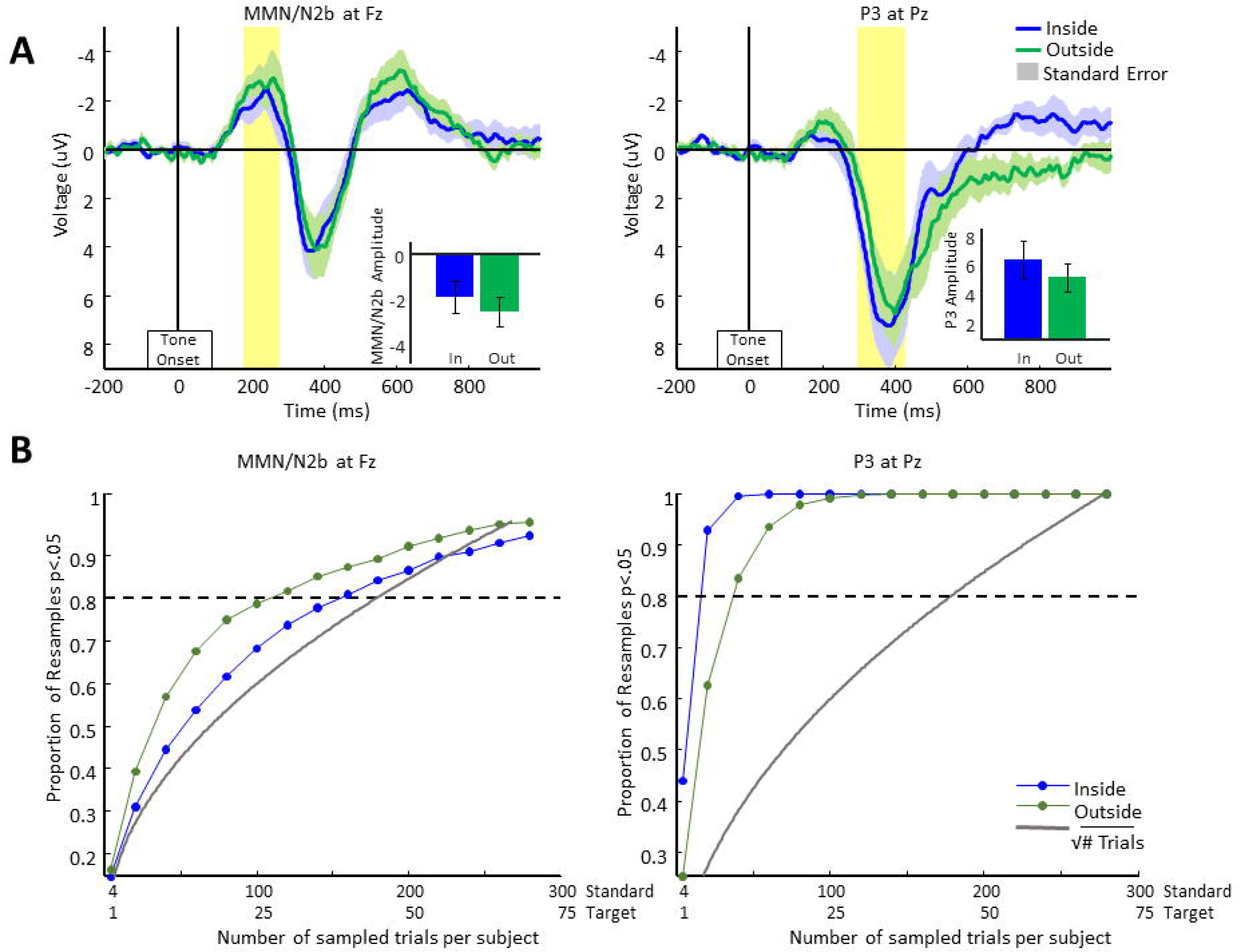
Data noise levels. A: Raw EEG data (with online band-pass and notch filters) for a representative subject for several minutes within each condition, shown at the Pz electrode location. B: Single-trial EEG spectra from electrodes Fz and Pz, computed with zero-padded FFTs on 504 auditory standard trial epochs of each subject, averaged over trials, then subjects. Shaded regions represent the standard error of the mean. C: Bar graph of average single-trial root mean square (RMS) grand average values collected during a 200 ms baseline period prior to tone onset, for 10,000 permutations of 360 randomly chosen standard and target trials for each subject. RMS values are averaged over all electrodes within each trial, then averaged over trials, and then averaged over subjects. Error bars represent standard deviation of the permuted distributions. D: Histogram of these 10,000 permuted single-trial RMS values. E: Bar graph of average ERP baseline RMS values, calculated using 10,000 randomly selected permutations of 360 standard and target trials for each subject. RMS of the baseline period is computed for each permutation, and the data averaged over trials. Error bars represent standard deviation of the permuted distributions. F: Histogram of these of these 10,000 permuted ERP baseline RMS values.

### 3.3. ERP Baseline Analysis

To test for noise that was not effectively averaged out over trials, we performed a similar analysis on noise levels within trial-averaged ERPs. To quantify the amount of noise in the participant average ERPs, we again used a permutation test of the RMS values in the baseline period. Complementary to the above single-trial analysis, this computation estimates the amount of phase-locked EEG noise in the data that is not averaged out over the trial with respect to onset of the tone. We randomly selected and averaged 360 standard trials without replacement from each participant’s standard non-artifact trials. The RMS values obtained were then averaged over EEG channels to create a grand average for all participants. Once each participant’s data were averaged together to compute second-order statistics, this made 10,000 permutations. The bar graph in Figure 2D depicts the means of these distributions, with error bars indicating the standard deviation of the distribution of permutation means. Figure 2E shows a histogram of the grand average RMS values calculated with these 10,000 permutations for each condition.Cycling outside (M_RMS-EEG_ = 0.4904; SD_RMS-EEG_ = 0.0181) produced a higher RMS value compared with the sitting inside condition (M_RMS-EEG_= 0.3685; SD_RMS-EEG_ = 0.0190; Wilcoxon rank sum test; z = −122.4709; p < 0.0001).

### 3.4. ERP Morphology and Topography

Figure 3A depicts grand-averaged ERPs following target and standard tones from electrode Pz, calculated from each participant’s corrected and artifact-free trials. The graph’s shaded regions depict the standard error of the mean for each time point. Similar levels of error are observed within each condition. Clearly, during outside cycling it is possible to measure ERPs with similar morphology and topography to those measured while sitting inside. Evident from the plots is the expected increase in amplitude for the P3 during target trials in posterior head locations (Pz), with a slightly greater P3 amplitude for targets in the inside condition.

An expected P3 oddball difference was demonstrated with increased positive voltages between 300-430 ms following the rare target tones, compared to the frequent standard tones. This time window was then used for all further P3 analyses. Additionally evident in Figure 4A was a difference in the target-standard difference in the time window of the MMN/N2b, with more negative voltage between 175-275 ms following rare target tones compared to the frequent standard tones. This time window was then used for all further analysis of the MMN/N2b, using electrode Fz where this was maximal.

**Figure 3.**
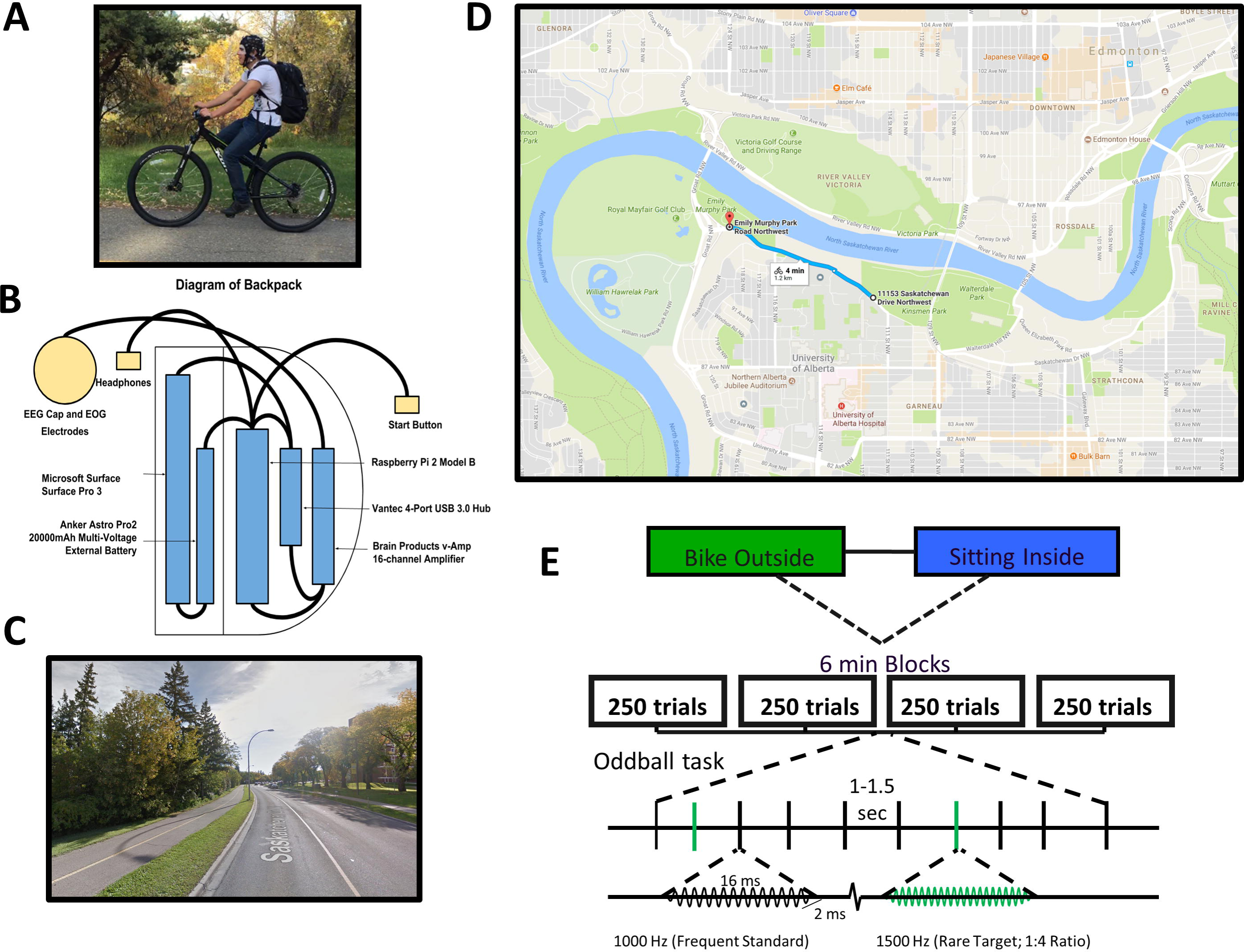
ERP grand averages. A: Grand-averaged ERPs computed at electrode Pz for all artifact-removed trials, corrected for eye movements, for both target (color) and standard (black) and tones. Positive is plotted down, and shaded areas depict standard error of the mean. B: Scalp topographies of the grand-averaged ERP difference between target and standard tones in the MMN/N2b and P3 time windows (indicated in yellow), 175–275 ms and 300–430 ms after the tone, respectively. EEG data was re-referenced after recording to the average of the left and right ear lobe electrodes. C: Difference wave ERPs from electrode Pz for both conditions, with shaded regions representing within-subject standard error of the mean for this difference, with between-subjects differences removed (Loftus & Masson, 1994). Regions highlighted in yellow depict the time window for MMN/N2b and P3 analysis as well as topographic plots.

Figure 3B illustrates topographies of these differences within the P3 and MMN/N2b windows. Topographies of the P3 time window show the expected posterior scalp distribution of activation for both conditions, while the MMN/N2b topographies reveal the expected distribution of activation toward the front of the head. Figure 3C plots the difference waves at Pz, which subtract the standard tone ERPs from their target tone ERPs for each subject. Shaded regions of this plot depict the within-participant standard error of the mean, with between-participant variation removed by the subtraction of standards from targets. This error estimate is therefore equivalent to that used in the t-test of this difference from zero (Loftus & Masson, 1994).

For both conditions there was a clear negative peak evident at approximately 220 ms at the Fz and even on the Pz electrode. A one-tailed, paired samples t-test comparing this MMN/N2b difference at electrode Fz averaged over the 175-275 ms time window centered around this peak revealed a significant MMN/N2b effect for the indoor condition (M_diff_= −1.9193; SD_in_= 2.4026; t(11) = −2.7674; p= 0.0092), as well as the outdoor cycling condition (M_diff_=-2.5988; SD_out_= 2.3145; t(11) = −3.8897; p=0.0013). Additionally as expected, a clear positive peak was observed at around 380 ms at Pz. A one-tailed, paired samples *t* test compared this P3 difference at electrode Pz averaged over the 300-430 ms time window revealed Significant P3 effect in both the indoor sitting (M_diff_= 6.0319; SD_in_= 4.4013; t(11) = 4.7474; p= 0.000301) and outdoor cycling conditions (M_diff_= 4.7865; SD_out_= 3.3884; t(11) = 4.8934; p= 2.3829e-04).

### 3.5. ERP Statistical Power

Figure 4A plots difference waves for both of the conditions for electrodes Pz and Fz. As shown, there is no significant difference of the MMN/N2b between the two conditions at Fz. A twotailed t-test between the inside sitting and outside cycling condition was performed to test for any condition differences between the targets-standards difference wave within the averaged 175-275 ms time-window, finding no significant difference (M_diff_= 0.6795; SD_dff_= 1.8018; t(11) = 1.3064; p= 0.2181).

**Figure 4.**
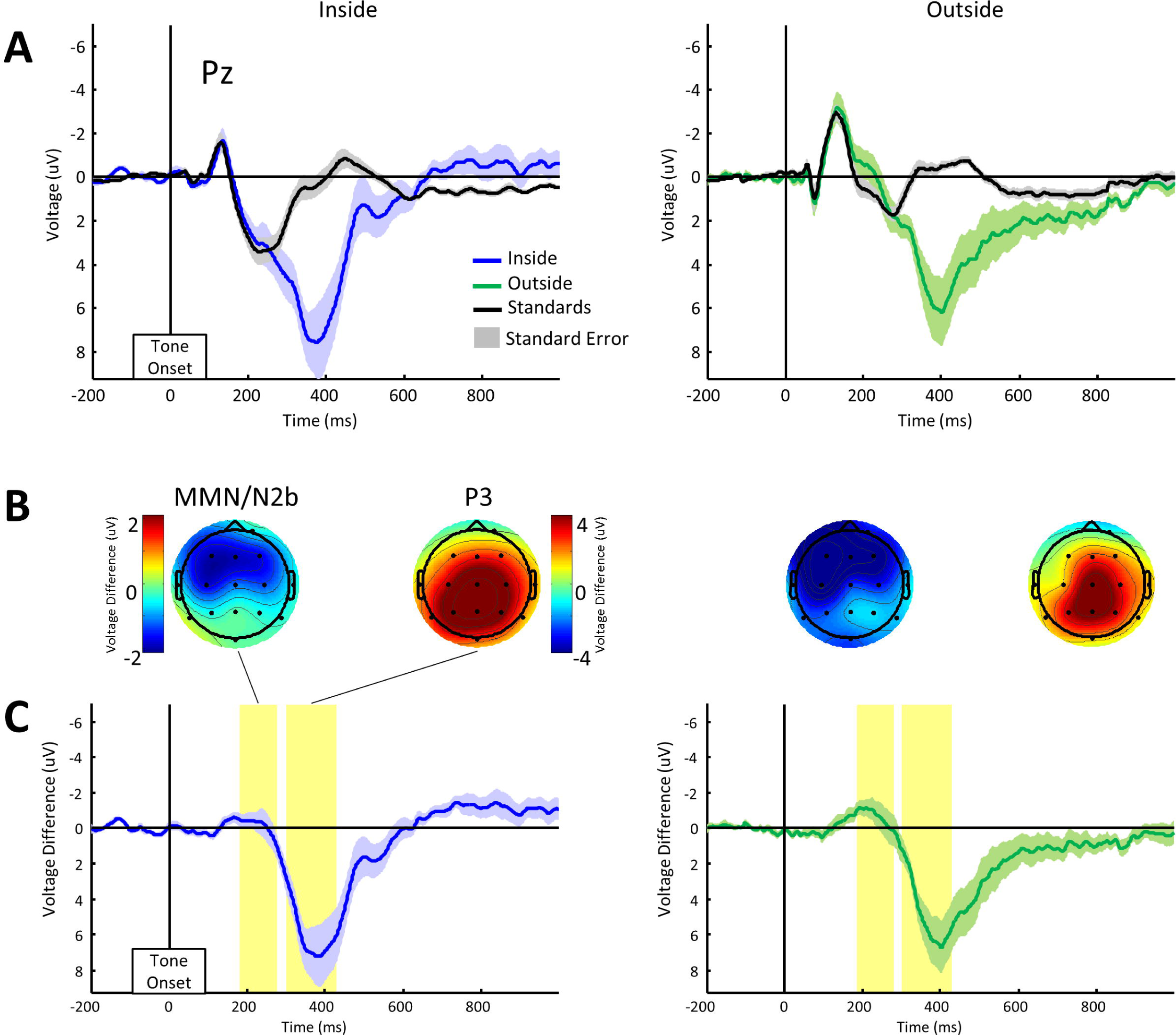
Difference waves and ERP power analysis. A: Difference waves indicating the average difference between target and standard trials for the two conditions are plotted for the Fz and Pz electrode locations. Yellow highlighted regions indicate main time windows compared, particularly the MMN/N2b at Fz (left) and the P3 at Pz (right). B: The results of a permutation test in which a number of trials selected within 10,000 permutations varied between 5 and 280, while keeping the 1:4 ratio of targets to standards. Randomly selected trials are averaged to calculate subject ERPs within each permutation for each number of trials. Differences in the P3 and MMN/N2b time window between standard and target trials is computed and compared using an across-subjects (paired samples) one-tailed t test (a= .05), before grand average statistics are computed. For each number of trials, the graph plots the proportions of the 10,000 permutations in which an uncorrected significant difference was obtained, for both conditions. The dashed horizontal line at 0.8 depicts the threshold to achieve 80% power to find an existing effect. The gray line represents a square root of the number of standard trials, scaled to a range between 0 and 1 on the vertical axis by dividing by the square root of the maximum number of trials.

To understand the different contribution to this effect between the MMN and N2b (Scanlon et al., 2017), we separated this window and differentially analyzed averages of an early time period for the MMN (100-200ms) and a later time period for N2b (200-300ms). No significant differences between conditions were found in the targets minus standards data when the two-tailed t-test was applied to either the MMN window (at Fz: M_diff_= 0.2770; SD_in_ =0.9285;SD_out_ = 1.1425;SDdiff=1.1643; t(11) =0.8242; p=0.4273; at Pz: Mf 0.2181; SDin = 0.9587;SD_out_ = 1.2823;SD_diff_=1.6279; t(11) =0.4641; p= 0.6516) or the later N2b window (at Fz: M_diff_= 0.8028; SD_in_ = 2.6399;SD_out_ =2.9497; SD_diff_=2.1544; t(11) =1.2908; p=0.2232; at Pz: M_diff_= 0.7560; SD_in_ =2.0251; SD_out_ = 2.3976; SDdiff=1.8850; t(11) = 1.3894; p=0.1922). Both the MMN and N2b were likely elicited here; however the current design does not allow us to disentangle these components from one other. From here on, we will therefore focus on the time window of a combined MMN/N2b between 175-275ms.

In figure 4A, the difference wave for the P3 at Pz does not appear to show any large differences between conditions. However a two-tailed *t* test performed to test for a significant difference in the averaged P3 difference (targets - standards) between the two conditions revealed a significantly reduced P3 difference in the outdoor cycling compared to indoor condition (M _in-out diff_=1.2454; SD_diff_=1.8072; t(11) =2.3871; p = 0.0360). Figure 3B shows topographies corresponding to these differences for the MMN/N2b and P3. It appears this difference is driven by a slightly earlier ramping up of the P3 in the inside condition.

Following evidence of increased single-trial and trial-averaged noise in the outside cycling condition compared to the indoor condition, one may expect to observe lower statistical power in the cycling condition. To test this prediction, we used a permutation procedure in which the 1:4 ratio of target to standard trials was held constant while varying the number of these trials contributing to the ERP average. Trial numbers increased from 1 target and 4 standard trial, by 20 standard trials, up to 300 standards and 75 target trials. We then randomly selected this number of trials from each participant’s overall trials, with replacement, and averaged over subjects to obtain second-order statistics (e.g. grand averages). This random replacement procedure was performed separately for each biking condition and for both the MMN/N2b and P3 analyses. 10,000 permutations of this procedure were done for each number of trials. Note that this procedure is unable to consider possible changes in ERP morphology over time in the task due to attention or habituation, and assumes that these influences are observed consistently across conditions and stimuli (Scanlon et al., 2017).

For each permutation, the single trials selected were averaged together to create separate participant ERPs for standard and target tones. Differences between target and standard tones were then calculated at electrode Fz between 175-275 ms and electrode Pz between 300-430 ms in order to measure MMN/N2b and P3 average values, respectively. We then compared these participant-average ERP differences using a paired samples *t*-test (*df=11,* one-tailed, *α*=.05). Figure 4B plots the proportion of 10,000 permutations in which the p-value obtained passed the threshold of significance, as a function of the number of targets and standards selected for each permutation. It is evident from this plot that the MMN/N2b for the cycling outside condition reached significance for 80% permutations (80% power gray line) with fewer trials (30 target/120 standard trials) than the inside condition (40 target/160 standard trials). The effect is opposite in the P3 in which the P3 for the cycling condition reached significance for 80% of the permutations with more trials (10 target/40 standard trials) than the inside condition trials (5 target/20 standard trials).

### 3.6. N1 and P2 Amplitudes

In order to measure effects of biking outside on stimulus processing in general, figure 5A-B plot grand averaged ERPs comparing inside vs outside for each condition at the Fz and Pz electrode locations, respectively. A visual inspection of these plots reveals increased amplitude of the N1 component between 100 and 175 ms in the outside cycling condition for both electrodes Fz and Pz, within both target and standard trials. Two-tailed *t*-tests indicated a significantly larger average amplitude of the N1 for outside compared to inside conditions at electrode Fz for both standard (M_diff_= 1.8694; SD_in_ =2.0195; SD_out_ = 1.5750; SD_diff_ = 1.2621; t(11) = 5.1309; p = 3.2783e-04) and target (M_diff_ 1.9978; SD_in_ = 2.1191; SD_out_ = 2.3810; SD_diff_ 1.6322; t(11) =4.2401; p = 0014) trials. Similar results were found for electrode Pz, indicating a significantly larger average N1 amplitude for the outside cycling condition, again within both standard (M_diff_ 1.5633; SD_in_ = 1.1053; SD_out_ = 1.0596;SD_diff_ = 0.7939; t(11) = 6.8210; p = 2.8724e-05) and target (M_diff_ 1.6707; SD_in_ = 1.4185; SD_out_ = 2.1360;SD_diff_= 1.6819; t(11) = 3.4411; p =0.0055) trials. The topography of this difference is shown in Figure 5C indicating a fronto-central distribution.

Visual inspection of Figure 5A-B also reveals decreased amplitude of the P2 component averaged between 175 and 275 ms, within both targets and standards of the Fz and Pz electrodes. Two-tailed t-tests revealed a significantly reduced P2 for the outdoor cycling condition at electrode Fz, mutually for standard (M_diff_= 2.3359; SD_in_ = 1.3812; SD_out_ = 1.4272; SD_diff_=1.3955; t(11) = 5.7986; p = 1.1953e-04) and target trials (M_diff_ 3.0154; SD_in_ = 3.2177;SD_out_ = 3.0679;SD_diff_= 2.4739; t(11) = 4.2223; p = 0.0014). This effect was also shown in the Pz electrode, with a significantly reduced P2 component for the outdoor cycling condition within both standard (M_diff_ 2.1640; SD_in_ = 1.6686;SD_out_ = 1.1310; SD_diff_ 1.2633; t(11) = 5.9338; p = 9.8189e-05) and target (M_diff_ 2.7335; SD_in_ = 2.3643;SD_out_ = 2.5763;SD_diff_= 1.9813; t(11) = 4.7792; p = 5.7211e-04) trials. The topography of this difference is shown in Figure 5C indicating a slightly more anterior fronto-central distribution.

**Figure 5.**
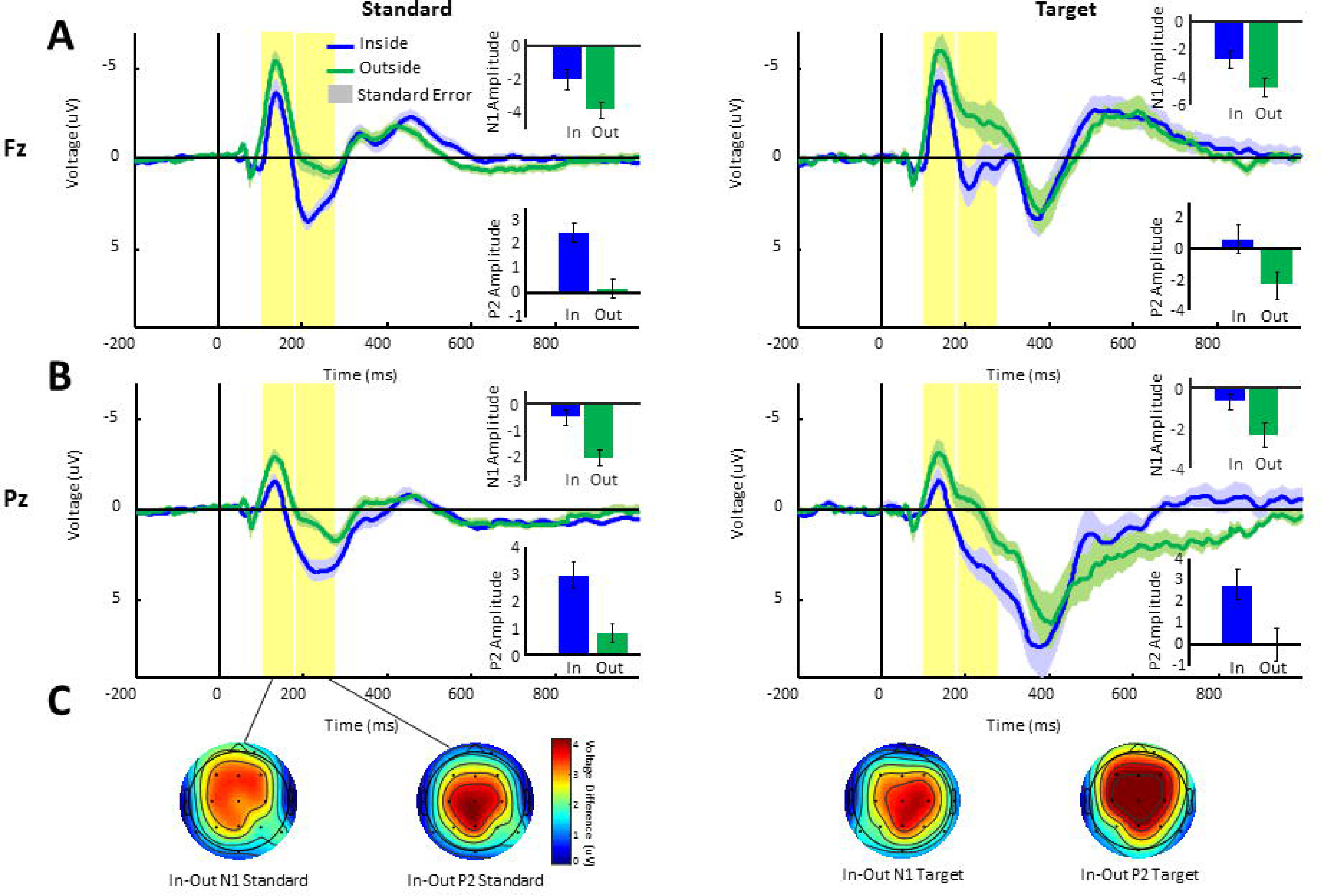
Grand average ERPs. A: Grand-averaged ERPs for the Fz electrode location, plotted separately to compare targets and standards between conditions. Shaded regions depict the standard error of the mean. Inset bar-graphs show the mean and standard-error across participants in the P2 window. B: Grand-averaged ERPs for the Pz electrode location, plotted separately to compare targets and standards between conditions. Shaded regions depict the standard error of the mean. C: Topographies of the N1 and P2 time window plotted for both inside and outside conditions.

## 4. Discussion

Here we assembled all the necessary components of an ERP experiment into a portable backpack and were able to acquire research quality ERP data in an auditory oddball task while users were riding a bicycle outdoors. We took advantage of recent advances in miniaturization of stimulus presentation devices (Kuziek, Shein, Mathewson, 2017) by using a raspberry Pi computer to present the auditory oddball task in headphones and to mark the EEG data for stimulus onsets. We then used a tablet computer to acquire marked EEG data from a Brain Products v-AMP 16-channel amplifier as the EEG system. Based on previous research on the noise levels of various electrode types (Mathewson, Harrison, and Kizuk, 2017), we used actively amplified electrodes with electrolyte gel. In this experiment, we had participants aim to not move their head side to side, and to avoid excessive eye movements. In future research we aim to minimize the need for these constraints further and allow for free viewing of the scene while biking. In past research on ERPs during video game play this has still allowed for reliable ERP measurement (Maclin et al., 2011; Mathewson et al., 2012).

As expected, we found that biking outside produced a great deal more noise in the EEG compared to sitting in the lab. This can be seen in Figure 2 as increased RMS in the baseline period of the EEG for both single trial data and the trial averaged ERP for each subject. For single trial data this increase in noise is about double the increase in RMS noise observed by individual pedalling inside the lab on a stationary bike compared to not pedaling (Scanlon et al., 2017). We attribute this additional noise to head and eye movements, and additional stimuli outside on the trail. The biking related increase in RMS noise we observed in the trial averaged ERP data (Figure 2E) is of similar magnitude as that found inside the lab in the previous study (Scanlon et al., 2017), suggesting that a great deal of this increased noise is not stimulus locked and is mitigated by trial averaging. Interestingly, when we considered the power spectra of this pre-stimulus noise, the results do differ from that observed in Scanlon and colleagues (2017) previous indoor biking study. While we observed the same increase in high frequency (beta band) noise when biking outside compared to sitting, we also observed an increase in low frequency noise in the delta (1-4Hz) range, as well as a very curious decrease in the peak of alpha normally observed in human EEG data. This decrease in the normally very prominent alpha peak indicates that the brain’s electrical activity when outside in a realistic setting is very different from the classic observation of a large alpha peak observed for almost a century (Berger, 1929). Given the understanding of alpha power as a measure of a pulsating inhibition in the brain (Mathewson et al., 2011), we suggest that this decrease in alpha observed outside is related to the bombardment of our senses of the multitude of visual, auditory, and other sensory information.

That we were able to measure P3 response to oddball stimuli when participants were riding outside on a bike, which were similar in magnitude to those measured inside the lab, is an important advance. Previously we had shown that in a video game inside the lab it was possible to measure research quality P3 response (Maclin et al., 2011). It has also been shown that reliable P3 response can be measured while individuals were walking outside (Debener et al., 2012; de Vos et al., 2014) as well as while biking outside (Zink et al., 2016). This is the first study to our knowledge that reliably compares the P3 observed between outside cycling and indoor laboratory conditions. As seen in Figure 3, the MMN/N2b and the P3 measured had similar topographies and morphologies inside the lab sitting compared to the same individuals outside the lab riding a bike.

In our analysis of the power of the ERP analysis as a function of the number of resampled trials used, we observed a moderate modulation of the ability to measure significant P3 and MMN/N2b effects. The MMN/N2b showed increased power, curiously, in the outside condition, with 50 fewer standard and 12 fewer target trials needed to allow for significant results on more than 80% of the resamples. For the P3, power to measure reliable differences between targets and standards we diminished in the outside condition, such that above 25 standard and 6 target trials more per person were needed. Our results indicate that to measure a reliable P3 outside with 100% power, around 125 artifact free standard and target trials would be needed, much less than then 1000 trials we recorded in this study.

As expected, a significantly reduced P3 was demonstrated while participants were cycling outside. This is in agreement with previous studies which have indicated that a lower amplitude P3 will be demonstrated when one’s cognitive resources are being divided between a main task and concurrent task (Kramer & Strayer, 1988; Polich, 1987; Polich & Kok., 1995; Wicken, Kramer, Vanesse, & Donchin, 1983), as well as with previous mobile EEG studies which showed a reduced P3 while cycling (Zink et al., 2016) and walking (Debener et al. 2012; de Vos et al., 2014).

The most interesting and unexpected finding of our results was something that we did not predict. However, there was a large difference in the magnitude of early ERP components observed when we compared directly the ERPs evoked by stimuli types when they were heard inside the lab compared to outside the lab on a bike. From around 100-300 ms, there is a large difference in the ERP such that the N1 peak is larger outside, and afterwards the P2 peak is almost non-existent. These differences have a fronto-central topography on the head. These changes in stimulus processing also do not seem to carry forward into differences in oddball detection, given the lack of differences in the P3 outside vs inside. Furthermore, when Scanlon and colleagues (2017) compared stationary biking in the lab vs. no movement, they did not find any modulations in the N1/P2, indicating that this effect is not due to the movement of biking itself.

The P2 has been associated with memory performance, working memory and semantic processing during tasks which are contextually based (Dunn, Dunn, Languis, & Andrews, 1998; Federmeier & Kutas, 2002; Lefebvre, Marchand, Eskes, & Connolly, 2005), while the N1-P2 complex is thought to reflect pre-attentive sound processing in the auditory cortex (Naatanen & Picton, 1987). The components of the N1-P2 complex, which include the P1, N1 and P2 appear to reflect several processes that overlap temporally and originate within or near the primary auditory cortex (Naatanen & Picton, 1987; Wolpaw & Pentry, 1975; Wood & Wolpaw, 1982). In particular, the auditory P2 component also appears to reflect the subjective difficulty involved in stimulus discrimination, as the P2 as well as the N1-P2 complex have both been reliably found to increase in amplitude following discrimination training (Atienza, Cantero, Dominguez-Marin, 2002; Hayes, Warrier, Nicol, Zecker, Kraus, 2003; Reinke, He, Wang, Alain, 2003; Trainor, Shahin, Roberts, 2003; Tremblay, Kraus, Carrell, McGee, 1997; Tremblay, Kraus, 2002; Tremblay, Kraus, McGee, Ponton, & Otis, 2001). The P2 is also believed to reflect an aspect of perceptual processing and top-down cognition, and has been hypothesized to represent a process of inhibiting the perception of repetitive stimuli which has been deemed unimportant to the task (Freunberger, Klimesch, Doppelmayr, & Höller, 2007; Luck & Hillyard, 1994). Additionally, the P2 has been proposed to be related to a process of suppressing irrelevant stimuli to allow discrimination between stimuli (Getzmann, Golob & Wascher, 2016; Kim, Kim, Yoon & Jung, 2008; Potts, 2004; Potts, Liotti, Tucker & Posner, 1996). The most relevant example of this was shown by Getzmann, Golob & Wascher (2016), demonstrating decreases in auditory P2 amplitude during speech discrimination while the participant ignored distracting background speech in a ‘cocktail party’ task, compared to a condition with no background stimuli.

We speculate that the modulation in the N1 and P2 we observed here is due to changes in discrimination difficulty and stimulus filtering outside. Our bike path was alongside a 50 km/hr road, which creates a great deal of auditory and visual noise unrelated to the task. The peak of the auditory spectra for sounds of traffic overlap with the 1000 and 1500 Hz stimuli used in the current study. Additionally, our task differed from most previous studies of the auditory N1 and P2, in that our background noises were cars passing by at relatively high speeds. While being irrelevant to the task, these stimuli may still be highly salient as they represent a real-life possible hazard to the individual, which may help to explain why this exact pattern of results has not been seen in previous similar studies. In follow up research in the lab we are testing these predictions by playing various background noise and visual noise while participants perform auditory and visual oddball tasks. It is possible that the differences in the N1, P2 and P3 we observe here are related to the differences in RMS noise from artifacts in the bike condition, which we remove only the largest of due to our very lenient rejection thresholds. We are however confident in the quality of the data in the bike condition considering the similarity of the ERP with that obtained on the same subjects while sitting still on a chair inside the lab. If artifacts decreased the P3 and P2 outside on the bike, it is also likely that they would decrease all the other components as well. This was not the case however, as the N1 was shown to increase, while other components were unchanged. In follow up studies inside the lab we have since replicated these N1 and P2 effects using recordings of traffic sounds from outside. Inside the lab we use stricter artifact rejection criteria and require participants to sit still on a chair, therefore we do not believe the N1 and P2 differences are due to motion artifacts.

Another unexpected finding was in the power analysis of the MMN, in which the outside condition appeared to reach 80% significant permutations with a smaller number of trials than the inside condition(Figure 4B), despite having a larger amount of noise by all other measures as well as no differences in mean amplitude. While it is difficult to speculate as to why this may have been, we believe that this may be due to noise from a few individual subjects. The standard deviation of the average MMN targets-standards difference was slightly higher for the inside condition (SD = 2.40) than the outside condition (SD = 2.31). The inside condition also had a wider range of values (Range_inside_= −7.3142-0.9951; Range_outside_ = −7.6030- −0.5067), three of which were greater than zero, while none of these values exceeded zero in the outside condition.

In this study we had participants ride as slow as comfortable to minimize noise from movement and to allow for sufficient recording time on the short trail we were having them ride on. Therefore, we don’t believe that any of our effects on ERP’s and data noise were due to effects of exercise on the brain. In a previous study inside the lab on stationary bikes (Scanlon et al., 2017), we did not observe these same modulations in the amplitude of the N1 and P2 when participants were pedalling a bike at a similar slow speed compared to sitting still on the bike. For the same reasons we don’t believe that the modulations in the N1 and P2 we observed are due to increases in noise due to movement, respiration or perspiration, given the slow speed of biking and the very similar noise levels compared to studies of indoor biking. We believe that any effects of acute exercise on brain processing would require a much higher intensity of biking (see Pontifex & Hillman, 2007; Yagi, Coburn &, Estes, Arruda, 1999; Grego et al., 2004; Bullock et al., 2017). In follow up research we plan to push individuals to faster biking paces and investigate changes in acute behavioural effects, ERP measurement ability, and measured noise in the data.

Cycling was used in this study for several methodological reasons. Firstly, for both the present study and the previous study with stationary cycling (Scanlon et al., 2017), as well as for our current set-up which includes wires connecting the electrodes and amplifier, cycling allowed a method to record EEG data during movement while minimizing the artifacts introduced into the EEG data. For example, when individuals walk or run, their head often moves up and down as they step, allowing wire movements and artifacts, while cycling often allows one to move forward in a relatively smooth straight-forward movement. The second reason is that cycling is a common activity which requires balance, steering control, and attention to one’s environment to be done correctly, making this a rich and unique activity to be explored experimentally. A simple oddball task was used to assess attentional allocation differences between sitting still inside and cycling outside in this study, which allowed us to infer that cycling at a slow and subaerobic pace outside is enough to alter the attentional resources allocated to the oddball task.

The limitations of our experimental setup continue to be in the bulk of equipment needed in the backpack. We have been using a large tablet computer to acquire the EEG data. To complement the raspberry Pi used for stimulus presentation, we have now started testing the use of a similar miniature computer (Latte Panda) that runs windows OS and can be used to record EEG data with brain vision recorder software (Brain Products, 2014). Further, our EEG amplifier itself is quite bulky, with electrode leads plugged in directly. Recent advances in both research grade EEG portable systems (Debener et al., 2015; Zink et al., 2016), would further improve the portability of such a system. It is also not known how well the active electrodes that we used in this study preform in movement situations, and some research has shown that data of similar quality can be measured with electrodes with no active amplification (Debener et al., 2012; de Vos, M., & Debener, S., 2014; Zink et al., 2016). Finally, consumer grade EEG hardware is becoming increasingly sophisticated, and very affordable Bluetooth EEG systems have been shown to measure research quality EEG and ERP data (Krigolson, Williams, Norton, Hassall, & Colino, 2017; Hashemi et al., 2016). We are also testing how these systems can be integrated into our experimental setup to further increase portability and minimize obtrusiveness of the equipment.

The ultimate goal of this research program will be to measure ERP’s to event that are naturally occurring out in the world, in order to gain a deeper understanding of how the brain works in everyday situations. This will require video capture and coding to identify moments in the EEG data when events of interest occur in the world. We have taken steps in this direction in the lab by testing the effectiveness of using virtual reality (VR) for stimulus presentation in ERP experiments. Further, we have begun filming our outdoor bike EEG sessions, and yoking the video recording to the EEG recording. We aim to start by using confederate actors to stimulate real life events (pedestrians, signals, obstructions, sounds), and move to video coding of real-world biking data.

### 4.1. Conclusion

In sum, here we have shown the ability to package an experimental EEG setup into a backpack and have individuals ride bicycles outside a path while engaged in an auditory oddball task. We reliably measured ERPs and can estimate the number of trials needed to measure reliable P3 responses outside on a bike. As expected, we found that while the P3 response was modulated by biking outside compared to sitting in the lab. However there was also a large modulation in the N1/P2 response to both standards and targets. We suggest that this modulation may be an effect of increased auditory filtering needed to focus on the task while ignoring the traffic sounds and sights.

## Acknowledgements

The authors declare no competing financial interests. This work was supported by a NSERC discovery grant and start-up funds from the Faculty of Science of KEM. The authors would like to thank Sayeed Kizuk for assistance with data collection. Reprints: Kyle E. Mathewson

